# The role of TGFβ signaling in Gli1+ tendon and enthesis cells

**DOI:** 10.1101/2022.03.03.482926

**Authors:** Lee Song, Mikhail Golman, Adam C. Abraham, Elazar Zelzer, Stavros Thomopoulos

## Abstract

The development of musculoskeletal tissues such as tendon, enthesis, and bone relies on proliferation and differentiation of mesenchymal progenitor cells. Gli1+ cells have been described as putative stem cells in several tissues and are presumed to play critical roles in tissue formation and maintenance. For example, the enthesis, a fibrocartilage tissue that connects tendon to bone, is mineralized postnatally by a pool of Gli1+ progenitor cells. These cells are regulated by hedgehog signaling, but it is unclear if TGFβ signaling, necessary for tenogenesis, also plays a role in their behavior. To examine the role of TGFβ signaling in Gli1+ cell function, the receptor for TGFβ, TbR2, was deleted in Gli1-lineage cells in mice at P5. Decreased TGFβ signaling in these cells led to defects in enthesis formation by P56, including deficient bone morphometry underlying the enthesis and decreased mechanical properties. *In vitro* experiments using Gli1+ cells isolated from mouse tail tendons demonstrated that TGFβ controls cell proliferation and differentiation through canonical and non-canonical pathways and that TGFβ directly controls the tendon transcription factor scleraxis by binding to its distant enhancer. These results have implications in the development of treatments for tendon and enthesis pathologies.

## INTRODUCTION

The musculoskeletal system is formed and maintained by a wide range of mesenchymal cells. Cells positive for Gli1, an essential hedgehog signaling transcription factor, have been described as putative stem cells in several tissues; a critical role for Gli1+ cells in development and response to injury has been shown for tendon, enthesis, teeth, bone, bone marrow, skin, colon, kidney, and heart [1-11]. These cells and their progenies have been localized to the tendon enthesis throughout postnatal development, eventually populating the entire fibrocartilage region between tendon and bone [12]. Furthermore, ablation of Gli1+ cells postnatally leads to a loss of mineralized fibrocartilage, suggesting that these cells play a critical role in enthesis mineralization [12]. Similarly, during tooth development, Gli1+ cells proliferate and differentiate into various cell types required for periodontal ligament and dental pulp formation [7, 8]. In bone, Gli1+ cells differentiate into various cell types at the growth plate during development and in the callus during fracture healing [10]. In tissues such as muscle, skin, and colon, Gli1+ cells have been reported to serve as stem cells within tissue-specific niches and actively regenerate injured tissues [1, 2, 13]. The regulation of these cells leading to their proliferation and/or differentiation is not fully understood, particularly for musculoskeletal tissues such as tendon and the enthesis.

Transforming growth factor beta (TGFβ) is a pleiotropic cytokine that plays important roles in several cellular functions, including proliferation, differentiation, and extracellular matrix (ECM) synthesis [14]. The TGFβ family of ligands works through the TGFβ receptors 1 and 2. Specifically, for the canonical pathway, a TGFβ ligand binds to the TGFβ receptor 2 (TbR2) and initiates a signaling cascade through Smad2/3. Alternatively, the noncanonical pathway results in activation of other transcription factors such as mitogen activated protein kinase (MAPK) and Akt [14, 15]. TGFβ signaling plays many critical roles in the formation of the musculoskeletal system. Deletion of TbR2 in Col2a-expressing cells resulted in smaller and disformed vertebral bones [16] and defects in the intervertebral disc (IVD) [17]; deletion in Col10a-expressing hypertrophic chondrocytes delayed bone collar calcification [18]; and deletion in osteocalcin-expressing osteoblasts caused increased trabecular bone mass and decreased cortical bone mass [19]. Deletion of TbR2 in Prx1-expressing cells also severely impaired the formation of tendons and ligaments [20]. Similarly, deletion of TbR2 in scleraxis (Scx) expressing tendon cells led to severe tendon disruption and eventually death by P14. Further studies revealed that TGFβ signaling was necessary to maintain tendon cell phenotype via Scx expression [21]. Recently, deletion of TbR2 in Gli1 expressing periodontal ligament (PDL) progenitors was shown to reduce PDL cell numbers [22]. These prior studies point to a crucial role for TGFβ in the formation of tendon and the mineralization of the tendon enthesis.

It is unknown if TGFβ signaling regulates the behavior of Gli1+ cells in tendon and the enthesis. To examine the role of TGFβ signaling in these cells, we performed a series of *in vivo* and *in vitro* studies. TbR2 was deleted in Gli1+ cells at P5 using Gli1Cre^ERT2^;TbR2^fl/fl^ mice. To study the effect of this deletion on the development of the enthesis, the phenotype of rotator cuff entheses was examined using biomechanical, morphological, and histological assays. To study the effect of TbR2 deletion on Gli1+ tendon cells, cells were isolated from tail tendon and their *in vitro* behavior in response to TGFβ and hedgehog agonist stimulation were examined. Results revealed that TGFβ signaling is critical for the formation of a functional tendon enthesis and controls Gli1-lineage cells through canonical and non-canonical pathways.

## MATERIALS AND METHODS

### *In Vivo* Studies

#### Animal models

To examine the effect of TGFβ signaling on tendon enthesis development, TbR2 was deleted in Gli1+ cells at P5. The following mouse models were purchased from The Jackson Laboratory (Bar Harbor, ME): *Gli1*^*tms(cre/ERT2)AIj*^/J (Gli1Cre^ERT2^, stock number 007913), B6;129-*Tgfbr2*^*tm1Karl*^/J (TbR2 floxed, stock number 012603), and B6.129(Cg)-*Gt(ROSA)26Sor*^*tm4(ACTB-tdTomato,-EGFP)Luo*^/J (mTmG, stock number 007676). To generate TbR2 deletion in Gli1-expressing cells, mice were crossed to produce Gli1Cre^ERT2^;TbR2^fl/fl^ conditional knockout (CKO) mice and Gli1Cre^ERT2^; TbR2^fl/wt^ control mice (control). In order to label Gli1-lineage cells, Gli1Cre^ERT2^;TbR2^fl/fl^ mice were mated with mTmG reporter mice to generate Gli1Cre^ERT2^;TbR2^fl/fl^;mTmG mice. In these mice, Gli1-expressing cells at the time of tamoxifen (TAM, Sigma-Aldrich, catalog number 06734) injection were labeled with membrane bound GFP while all the other cells were labeled with membrane bound td-Tomato [23]. To induce Cre^ER^ nuclear localization and delete the TbR2 allele, mice were subjected to subcutaneous injection of TAM. TAM was dissolved in sterile corn oil (Sigma-Aldrich, catalog number: C8267) at a concentration of 10 mg/ml and subcutaneously injected at 100 mg/kg body weight at P5 and P7 [12, 23]. Gli1Cre^ERT2^;TbR2^fl/fl^ mice were used at 8 weeks of age. Due to the limitation of available mice, the control group for bone and mechanical property included both Gli1-Cre^ERT2^;TbR2^fl/wt^ and TbR2^fl/fl^ mice since both strains express TbR2. All the mouse experiments were approved by the Institutional Animal Care and Use Committee (IACUC) of Columbia University. The phenotypes of the mice were determined using biomechanics, bone morphometry, and histologic analyses.

#### Micro computed tomography (micro-CT)

Mice were euthanized at P56, humerus-supraspinatus tendon/muscle samples were isolated, and samples were stored in saline-soaked gauze at -20 ºC until scanning (N=14 for control, N=11 for CKO). Specimens were thawed and micro-CT scanning was performed using a Skyscan 1271 scanner (Bruker Corporation) at an energy of 60 kV, intensity of 166 μA, 0.25 μm aluminum filter, and 5 μm resolution. Scanned images were evaluated using a custom segmentation algorithm to separate cortical and trabecular bones of the humeral head proximal to the growth plate (CTAn, Bruker Corporation). Bone mineral density (BMD) and total mineral density (TMD), bone volume fraction (BV/TV), trabecular number (Tb.N), trabecular thickness (Tb.Th), and trabecular spacing (Tb.Sp) were measured. Tendon cross-sectional area was measured by thresholding μCT images of sagittal slices through the tendon. The minimum cross-sectional area was used for mechanical property analysis [24, 25].

#### Biomechanical testing

After μCT scanning, the supraspinatus muscle was carefully scraped from the tendon, the humeral bone was mounted in a custom 3D printed fixture, and a uniaxial tensile test to failure was performed using an Electroforce 3230 testing frame (TA Instruments) (N=14 for control, N=11 for CKO) [26]. The testing protocol consisted of 5 cycles of preconditioning (2% strain, 0.2% /s), 180 s recovery, and extension to failure at 0.2% /s. Strain was determined from grip-to-grip displacement relative to the initial gauge length. Stress was determined using the minimum cross-sectional area of the tendon, as determined from micro-CT. Structural properties were determined from load/deformation curves, including failure load (maximum load), stiffness (slope of linear portion of load/deformation curve), and work to failure (area under load/deformation curve through yield). Material properties were determined from stress/strain curves, including strength (maximum stress), modulus (slope of linear portion of stress/strain curve), and resilience (area under the curve through yield).

#### Histology and immunohistochemistry

Mice were euthanized at P16 when entheses moralization occurs, humeral-supraspinatus tendon/muscle samples were isolated, and samples were fixed in 4% paraformaldehyde (ThermoFisher Scientific, catalog number 50-980-487) at 4ºC overnight (N=8 for control, N=3 for CKO). The specimens were then decalcified in 10% EDTA (pH 7.4) (Poly Biotech). Specimens were rinsed in PBS and dehydrated in 30% sucrose (Fisher Scientific, catalog number S5-500). The specimens in sucrose were then embedded in OCT compound (VWR, catalog number 25608-930) and sectioned into 9 μm-thick sections using a cryostat (Ag protect, CM1860, Leica). The sections were fixed in 10% neutral buffered formalin (StatLab, catalog number 28600-1) for 20 minutes, and permeabilized in 1% Triton X-100 (Sigma-Aldrich, catalog number T9284). After blocking with 5% bovine serum albumin (BSA, fraction V) (Fisher Scientific, catalog number BP1600), the sections were incubated with either 1:250 anti Ki67 (Abcam, catalog number ab15580), 1:100 anti-p-cJun (Cell Signaling Technologies, catalog number 3270), anti-SPARC (R&D Systems, catalog number: AF942-SP), or 1:100 biotinylated anti-collagen 1a (GeneTax, catalog number GTX36577) overnight at 4°C. The sections were then incubated with 1:1000 anti-rabbit IgG Alexa Fluor 555 (Invitrogen, product number A-27039), anti-mouse IgG Alexa Fluor 647 (Invitrogen, product number A-21235), or anti-goat IgG Alexa Fluor 555 (Invitrogen, catalog number A-21431) after washing 3 times with Tris Buffered Saline (Boston BioProducts, catalog number IBB-588) containing 0.5% TWEEN®-20 (Sigma-Aldrich, catalog number P1379-100ML) (TBST). The sections were then rinsed with TBST and mounted with VECTASHIELD Plus mounting media with DAPI (Vector Laboratory, reference number H-2000). The images were captured with either a Zeiss Observer 7 microscope/Axiocam 702 camera (for Alexa Fluor 555) or Nikon A1RMP Multiphoton Confocal Microscope (for Alexa Flour 647). Cells positive for antibodies and/or GFP were counted manually. To examine tendon and enthesis morphology, a separate group of sections were stained with Safranin O per manufacturer’s instruction (American MasterTech, reference number: KTSFO).

#### Flow cytometry

To analyze the number of Gli1-lineage cells in the supraspinatus, infraspinatus, and Achilles tendon entheses, tissues were dissected from Gli1-Cre^ERT2^;TbR2^fl/wt^;mTmG heterozygous control and Gli1-Cre^ERT2^;TbR2^fl/fl^;mTmG CKO mice. The pooled tissues were chopped into small pieces with a razor blade and digested in 5 ml of complete medium MEMα (ThermoFisher Scientific, catalog number 1256107) containing 10% fetal bovine serum (Gemini Bio-products, catalog number 900-108), 100 U/ml penicillin-streptomycin (ThermoFisher Scientific, catalog number 15140122) and 4-5 mg/ml type II collagenase (Worthington Biochemical Corp, catalog number 004117) at 37 ºC with 225 rpm shaking. After 1.5 hours, 5 ml of complete medium was added to the digestion. The digested tendon fragments were then passed through a 20 G needle to break any clumps and then filtered through a 70 μm cell strainer (Fisher Scientific, catalog number 08-771-2). The cells were then spun down at 350 xG for 5 min and washed once with 5 ml fresh complete medium. The cell suspensions were stained with CD45-Brilliant violet 711™ (1:100, Biolegend, catalog number: 103147) and DAPI (1:5,000, Invitrogen, catalog number: D1306). The stained cells were passed through a LSRII cell analyzer (BD Biosciences) and analyzed using FCS Express software. The percentage of positive cells for each population was presented after CD45+ hematopoietic cells and DAPI+ dead cells were excluded.

### *In vitro* studies

#### Cell culture

To examine the role of TGFβ signaling in Gli1-lineage cell function, Gli1-lineage cells labeled with GFP were isolated from tail tendons and cultured, as previously described [27]. Briefly, tail tendon fascicles were pulled from their attachments, chopped into small pieces with a razor blade, and digested in 5 ml of complete medium MEMα (ThermoFisher Scientific, catalog number 1256107) containing 10% fetal bovine serum (Gemini Bio-products, catalog number 900-108), 100 U/ml penicillin-streptomycin (ThermoFisher Scientific, catalog number 15140122) and 4-5 mg/ml type II collagenase (Worthington Biochemical Corp, catalog number 004117) at 37 ºC with 225 rpm shaking. After 1.5 hours, 5 ml of complete medium was added to the digestion. The digested tendon fragments were then passed through a 20 G needle to break any clumps and then filtered through a 70 μm cell strainer (Fisher Scientific, catalog number 08-771-2). The cells were then spun down at 350 xG for 5 min and washed once with 5 ml fresh complete medium. The cells were then re-suspended in 3 ml of complete medium and plated into a 35 mm cell culture plate until they reached 80% confluence. The cells were then lifted by trypsin/EDTA (ThermoFisher Scientific, catalog number 25200056) and re-plated at a 1:2 ratio.

The cultured cells were separated by flow cytometry (Influx, BD Bioscience) at passage 3 based on their expression of either GFP or td-Tomato and analyzed using FCS Express software. The sorted cells were then further expanded. The cells were then replated at 3,000 /cm^2^ and starved in MEMα with 1% FBS overnight. For Western blot analysis, the starved cells were then treated with 5 ng/ml of recombinant murine TGFβ 1 (PEPROTECH, catalog number AF-100-21C). Cell responses were evaluated at 0, 15, 30, 60, 120, and 1440 minutes. Cells were lysed with RIPA Lysis and Extraction Buffer (ThermoFisher Scientific, catalog number: 89900) supplemented with Halt™ Protease and Phosphatase Inhibitor Cocktail (ThermoFisher Scientific, catalog number 78440) according to manufacturer’s instructions. For gene expression analysis, the starved cells were treated with recombinant murine TGFβ 1 (5 ng/ml) or hedgehog agonist (Hh-Ag1.5, Xcess Bioscicences Inc., CAS# 612542-14-0, 0.1μM) for 1, 4, 24, 48, and 72 hours. Total RNA was extracted using RNeasy Mini Kit (Qiagen, catalog number:74104). For immunofluorescence, cells were grown on a cover slip and treated with recombinant TGFβ 1 (5 ng/ml) for 60 minutes.

#### Western blot analysis

Western blot analyses were performed on cultured Gli1+ cells to determine signaling pathways induced by TGFβ (N=5 cell isolations for control, N=3 cell isolations for CKO). Whole cell lysates were quantified with the Pierce™ BCA Protein Assay Kit (ThermoFisher Scientific, catalog number: 20227). 20 μg of total protein was separated on an 8% SDS-PAGE, transferred to a 0.2 μm nitrocellulose membrane (BioRad, catalog number 1620112), and probed with 1:1000 of anti-phospho-Smad2 (Cell Signaling Technology, catalog number 18338S), anti-Smad2 (Cell Signaling Technology, catalog number 5339S), anti phospho-cJun (Cell Signaling Techology, catalog number 3270S), anti cJun (Cell Signaling Technology, catalog number 9165S), anti-phospho-SAPK/JNK (Cell Signaling Technology, catalog number 4668S), anti JNK (Cell Signaling Technology, catalog number 9252S), anti-phospho-p38 MAPK (Cell Signaling Technology, catalog number 4511S), anti-p38 MAPK (Cell Signaling Technology, catalog number 8690S), anti-phospho-p44/p42 MAPK (pErk, Cell Signaling Technology, catalog number 8690S), anti-p44/p42 MAPK (Erk, Cell Signaling Technology, catalog number 4695S), and anti-GAPDH (Cell Signaling Technology, catalog number 5174S). The primary antibody was detected with an anti-rabbit IgG, HRP-linked secondary antibody (Cell Signaling Technology, catalog number 7074S; 1:4,000) and visualized with either SuperSignalTM West Femto Maximum Sensitivity Substrate (ThermoFisher Scientific, catalog number 34094) or SuperSignal™ West Pico PLUS Chemiluminescent Substrate (ThermoFisher Scientific, catalog number 34577) for phospho-JNK and GAPDH. Images were captured with an Azure 600C Imager (Azure Biosystems) and quantified using Azurespot software. Between application of different antibodies, the nitrocellulose membrane was stripped with Restore™ Plus Western Blot Stripping Buffer (ThermoFisher Scientific, catalog number 21059) according to manufacturer’s instructions.

#### Quantitative polymerase chain reaction (qPCR)

To determine expression patterns of cells induced with TGFβ or Hh Agonist, qPCR was performed on cultured Gli1+ cells (N=8 cell isolations for control, N=3 cell isolations for CKO). One hundred nanograms of total RNA were reversely transcribed into complementary DNA (cDNA) using Maxima First Strand cDNA Synthesis Kit for RT-qPCR (ThermoFisher Scientific, catalog number K1641). The cDNA was diluted 10-fold with water and 2 μL of diluted cDNA was quantified for each specific gene using Power SYBR™ Green PCR Master Mix (ThermoFIsher Scientific, catalog number 4368706) in a QuantStudio™6 Pro Real-Time PCR System (ThermoFisher Scientific, catalog number: A44288) according to manufacturer’s instructions. The primer pairs for each gene were designed using an online primer design tool (Primer3, http://primer3.ut.ee/). The primer sequences are listed in Table 1.

**Table 1.**
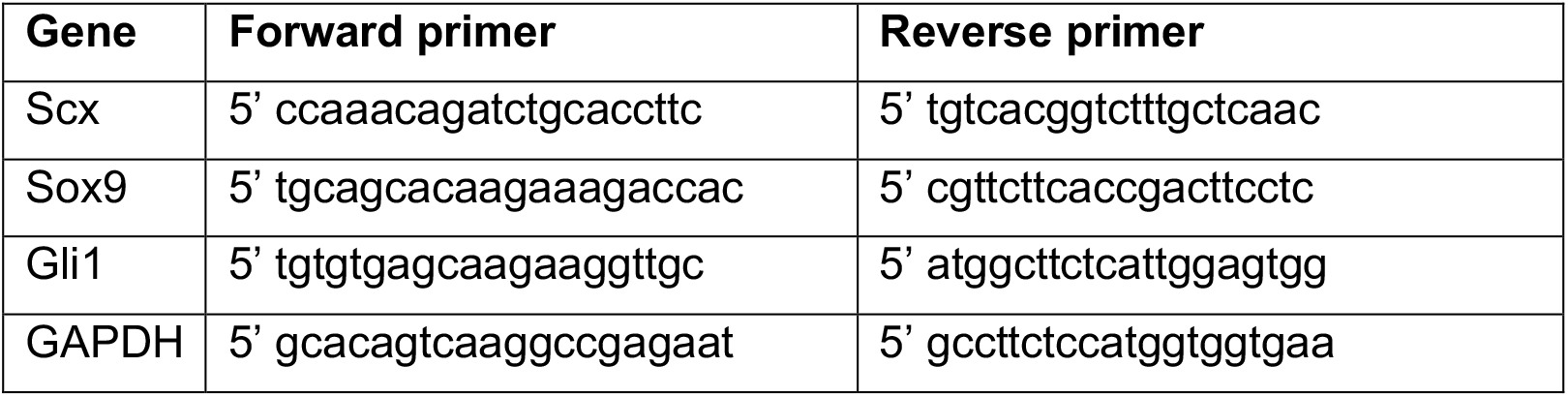
Primer sequences for qPCR.

##### Chromatin immunoprecipitation (ChIP)

ChIP analysis was performed to explore how TGFβ controls expression of Scx, a key tendon enthesis transcription factor [28] (N=2 cell isolations for control and N=2 cell isolations for CKO). Tail tendon fibroblasts from control and CKO mice were grown to 90% confluence in 15 cm cell culture plates. The cells were treated with TGFβ 1 (5ng/ml) for 1 hour up to 24 hours in MEMα-1. The chromatins were harvested using Cell Signaling Technology’s SimpleChIP^R^ Enzymatic Chromatin IP Kit (Magnetic Beads) and precipitated with anti-mouse Smad4 antibody (Cell Signaling, catalog number 46535, lot 2) according to manufacturer’s protocols (Cell Signaling, #9003). Briefly, tendon fibroblasts were seeded at 2×10^6^ per 15 cm plate in complete media. When the cells reached to 90-95% confluence, the media was switched to MEMα -1 overnight. The cells were treated with TGFβ 1 (5ng/ml) for 1 hour to 24 hours on the following day in 20 mL MEMα -1 or fresh 20 mL MEMα -1 for un-treated controls. At the end the stimulation, 540 μL of 37% formaldehyde (Fisher Scientific, catalog number BP531) were added to cross-link the proteins with DNA. The cross-linking was stopped by adding glycine. The cells were then washed with cold PBS and pelleted in 2 ml cold PBS at 2,000 x G for 5 min. The cell pellets were then digested with 0.3 μL of Micrococcal Nuclease for 20 min at 37 ºC. The digestion was stopped by adding 10 μL of 0.5 M EDTA. The digested chromatins were then fractioned by sonication via 3 × 30 s pulses with 20% Duty Cycle, Output Control setting at 3 in a Branson SONIFIER 250 sonicator (Branson Ultrasonics Corp). The chromatins were then split into two parts: 1 part was saved as an input control and 1 part was incubated with anti Smad4 rabbit monoclonal antibody at 1:100 overnight at 4 ºC. The following day, 30 μL of Protein G magnetic beads were added to the antibody-chromatin mix and incubated for 2 hours at 4 ºC. The magnetic beads-Smad4-chromatin complexes were then purified in a DynaMag2 magnetic particle concentrator (Invitrogen). The purified complexes were eluted in a high salt solution and reverse cross-linked by proteinase K. The immunoprecipitated chromatins and input control DNA were further purified with a DNA purification kit (Cell Signaling, catalog number 14209). The precipitated DNA was quantified using qPCR with specific primers for each potential binding site (Table 2). The binding capacity was calculated as the percentage of input using the formula: percentage of input = 2×2^(Ct of input-Ct of IP)^. To pool data from different experiments, the fold change between TGFβ treated samples was compared to untreated samples. Those samples were then examined for potential Smad4 consensus binding sites using ALGGEN-PROMO from 10 kb upstream and 10 kb downstream of mouse Scx gene transcription start site (TSS) [29] and compared with the conserved regulatory regions reported by Pryce et al [30].

**Table 2.**
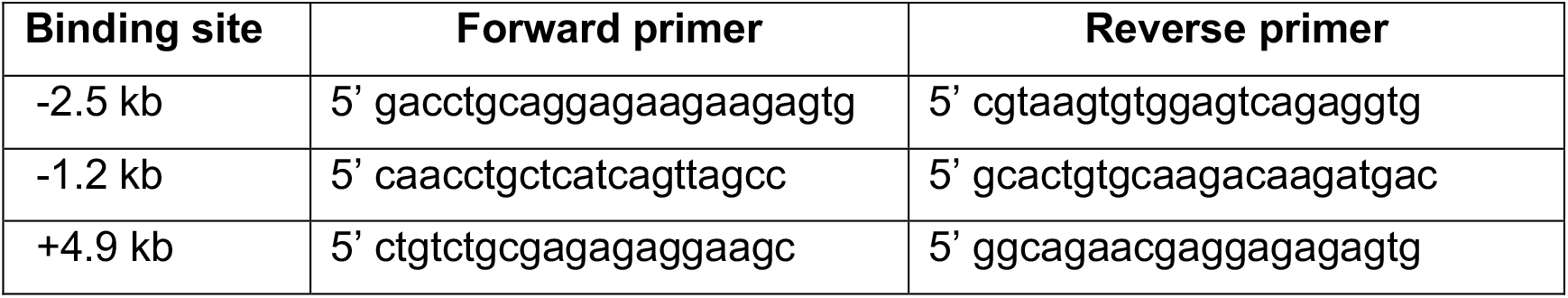
Primer sequences for ChIP.

#### Immunocytochemistry

To examine the effectiveness of TGFβ deletion in the mouse models, sorted cells from reporter mice were cultured and examined using immunofluorescence. After TGFβ treatment, tendon fibroblasts were washed 2 times with cold PBS and fixed in 0.4% formaldehyde (Fisher Scientific, catalog number BP531-500) for 20 min. The cells were then permeabilized in 1% Triton-X100 for 2 min. After three 5 min washes with TBST, the cells were blocked with 5% bovine serum albumin (Sigma-Aldrich, catalog number A3803-100G) for 30 minutes and 1:100 anti-Smad2 (Cell Signaling Technology) for 2 hours. After 3 washes with TBST, the cells were incubated with 1:1000 Donkey anti-Rabbit IgG (H+L) Highly Cross-Adsorbed Secondary Antibody, Alexa Fluor 488 (ThermoFisher Scientific, catalog number A21206) for td-Tomato positive cells or 1:1000 of Donkey anti-Rabbit IgG (H+L) Cross-Adsorbed Secondary Antibody, Alexa Fluor 555 (ThermoFisher Scientific, catalog number: A-31572) for 1 hr. The cover slips of cells were then mounted onto slides with VECTASHIELD® Antifade Mounting Media with DAPI (VECTOR Laboratories, catalog number: H-1200) after 5 TBST washes. Images were taken on a Nikon A1RMP Multiphoton Confocal Microscope.

### Statistics

All data are presented as mean plus/minus standard deviation. The statistical significance between control and CKO groups in FACS and histology data was determined by student t-test. The statistical significance for qPCR and western blots was decided with ANOVA using Prism software (GraphPad). The micro-CT and mechanical testing data was determined by ANCOVA to correct for gender effect using SPSS 26 (IBM).

## RESULTS

### TGFβ signaling was blocked in Gli1-lineage cells

To isolate Gli1-expressing cells whose TGFβ receptor 2 (TbR2) was deleted, Gli1Cre^ERT2^;TbR2^fl/fl^ mice were crossed to Rosa26R/mTmG reporter mice to generate Gli1Cre^ERT2^;TbR2^fl/fl^;mTmG mice. Tamoxifen (TAM) injection at P5 and P7 led to deletion of TbR2 and expression of GFP in Gli1-lineage cells (Figure 1A). Thus, TbR2-deleted cells expressed GFP and all other cells (that did not express Gli1) expressing td-tomato (Figure 1A). Tail-derived tendon fibroblasts (TFs) were expanded and separated based on their GFP (green fluorescence) and td-tomato (red fluorescence) expression in a flow cytometer and cultured. To test if TGFβ signaling pathway was blocked in Gli1Cre^ERT2^;TbR2^fl/fl^ cells, the sorted td-Tomato+ and GFP+ cells were treated with TGFβ 1 (5ng/ml) for 60 minutes and then stained for Smad2. In td-Tomato positive (i.e., wild type) cells, Smad2 protein was primarily localized to the nuclei (Figure 1B, bottom row). In contrast, Smad2 remained in the cytoplasm of TbR2-deleted GFP+ cells (Figure 1B, top row). Consistent with this result, Western blot analysis demonstrated marked decreases in pSmad2 in TbR2-deleted GFP+ cells compared to wild type control cells (Figure 1C). These data demonstrate that TGFβ signaling was effectively blocked in Gli1Cre^ERT2^;TbR2^fl/fl^ tendon cells.

**Figure 1.**
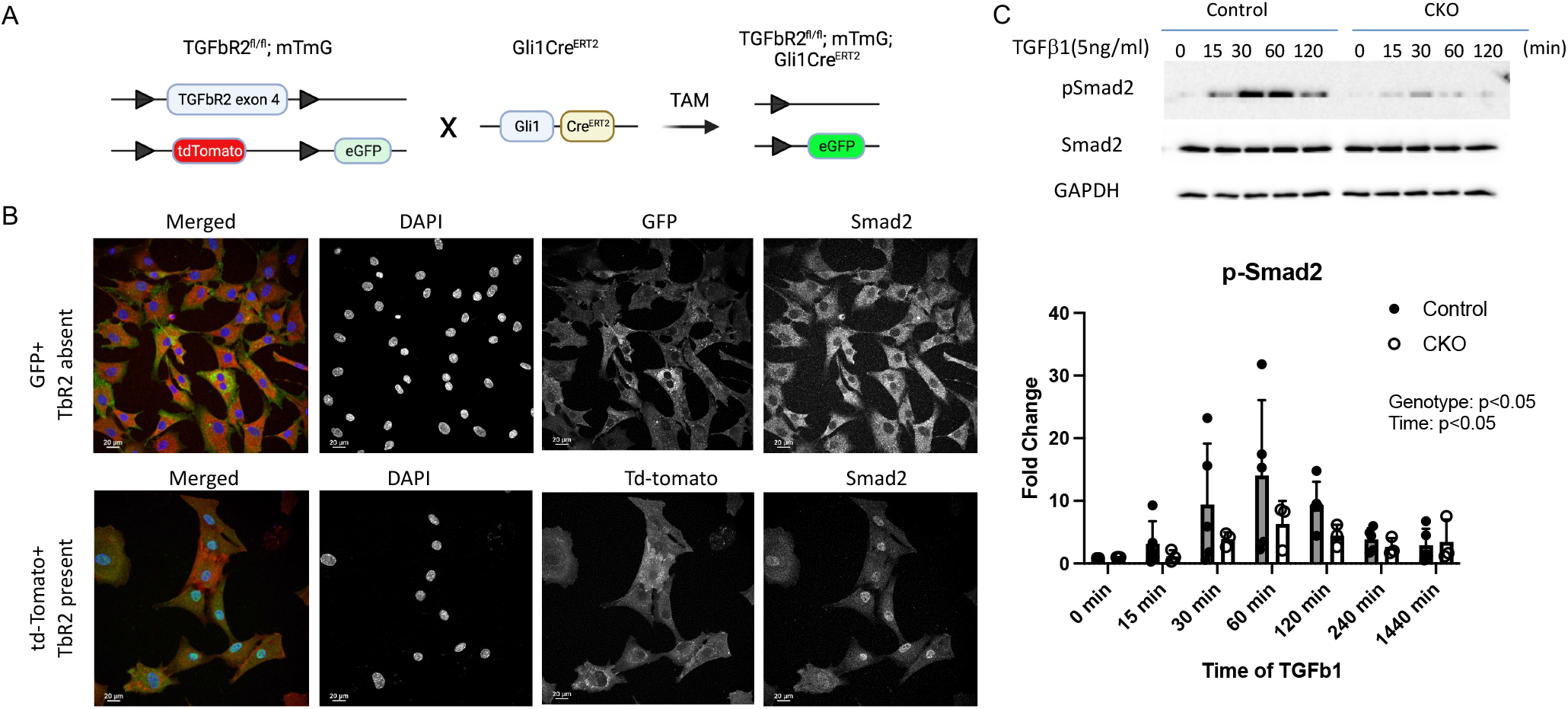
Generation and validation of Gli1Cre^ERT2^;TbR2^fl/fl^;mTmG mice. **(A)** The strategy for generating reporter mice whose Gli1-lineage TbR2-deleted cells were tagged with GFP. **(B)** Immunocytochemistry of cultured cells showed that TGFβ -induced Smad2 nuclear localization was blocked in Gli1-lineage (green) cells with TbR2 deletion (top row), in contrast to non-Gli1-lineage cells (red) (bottom row). **(C)** Western blot analysis of cultured cells demonstrated substantially reduced Smad2 phosphorylation in Gli1Cre^ERT2^;TbR2^fl/fl^ cells (CKO) cells compared to control cells.

### Postnatal deletion of TbR2 in Gli1-lineage cells led to mechanical and morphological defects in the enthesis

TbR2 was deleted in Gli1-lineage in the early post-natal period (Figure 2A). Uniaxial tensile testing of supraspinatus tendon entheses revealed that the Gli1Cre^ERT2^;TbR2^fl/fl^ (CKO) mice had lower maximum force, work to failure, and resilience compared to TbR2^fl/wt^ control (CTRL) mice (Figure 2B). Micro-CT analysis of humeral head bone near the supraspinatus tendon enthesis showed that, compared to CTRL mice, CKO mice had lower cortical bone thickness, smaller total cross-sectional area (TtAr), less bone volume to total volume ratio (BV/TV), lower bone mineral density of articular bone (trabecular BMD), and lower trabecular bone thickness (TbTh) (Figure 2C,E). Histologic appearance of the supraspinatus tendon enthesis at P16 (i.e., immediately after enthesis mineralization) was similar in the CKO mice compared to control mice, with some decreased staining for fibrocartilage (Figure 2D).

**Figure 2.**
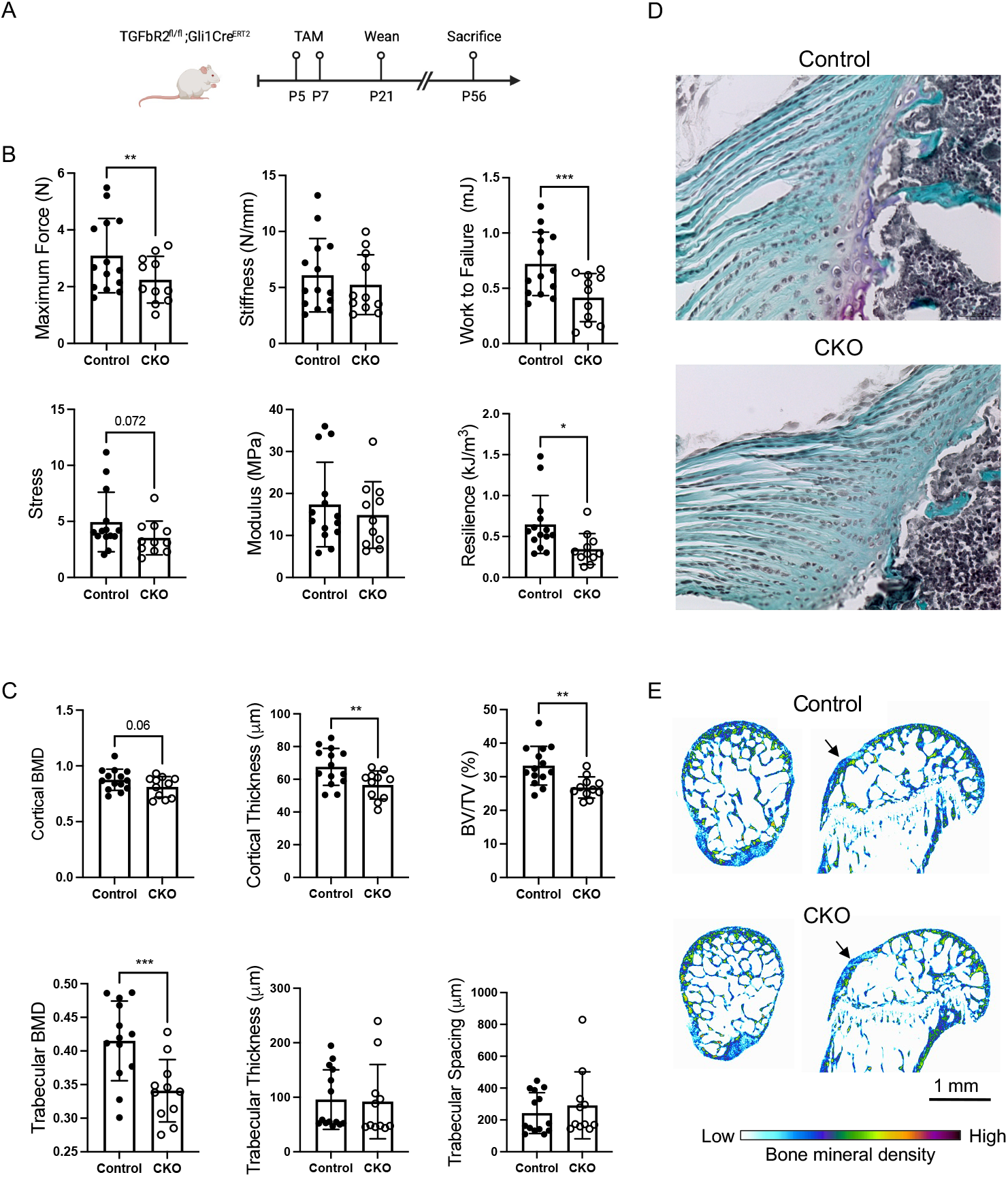
TbR2 deletion in Gli1-lineage cells led to reduced enthesis mechanical properties and altered bone morphometry. **(A)** Tamoxifen (TAM) was administrated at P5 and P7 to induce deletion of TbR2 in Gli1-expressing cells (CKO). **(B)** The mechanical properties of the supraspinatus tendon enthesis were reduced in CKO mice compared to CTRL mice. * p<0.05, ** p<0.01, *** p<0.001. **(C)** Deletion of TbR2 in Gli1-expressing cells led to altered bone morphometry of the humeral head cortical and trabecular bone. **(D)** Representative histologic sections for control and CKO supraspinatus tendon entheses (Safranan O stain, tendon is on the left and bone is on the right of the sections). **(E)** Representative microCT images show reduced trabecular bone and lower bone mineral density adjacent to the supraspinatus tendon enthesis (arrows). ** p<0.01, *** p<0.001.

### ECM changes in Gli1Cre^ERT2^;TbR2^fl/fl^ CKO mice were limited to Gli1-lineage cells

Collagen 1a (COL1a) production is an established consequence of TGFβ signaling in musculoskeletal tissues, as seen during tissue fibrosis and scar formation, including for tendon and enthesis [31-34]. Histomorphometric analysis revealed a trend towards increased numbers of COL1a+ cells in CKO entheses compared to Control entheses, however the percentage of COL1a+ cells was not different between the two groups (Figures 3A, S1A). When considering only Gli1-lineage (i.e., GFP+) cells, there were trends for decreased numbers and percentage of COL1a+ cells in CKO compared to Control enthesis (Figure 3A). Secreted protein acidic and rich in cysteine (SPARC, a.k.a osteonectin) is a major non-collagen ECM protein found in tendon and mineralized tissues [35, 36]. The numbers and percentage of SPARC+ cells were similar between CKO and Control entheses (Figures 3B, S1B). When considering only Gli1-lineage (i.e., GFP+) cells, there was a significant reduction in the percentage of SPARC+ cells in CKO compared to Control enthesis (Figure 3B).

**Figure 3.**
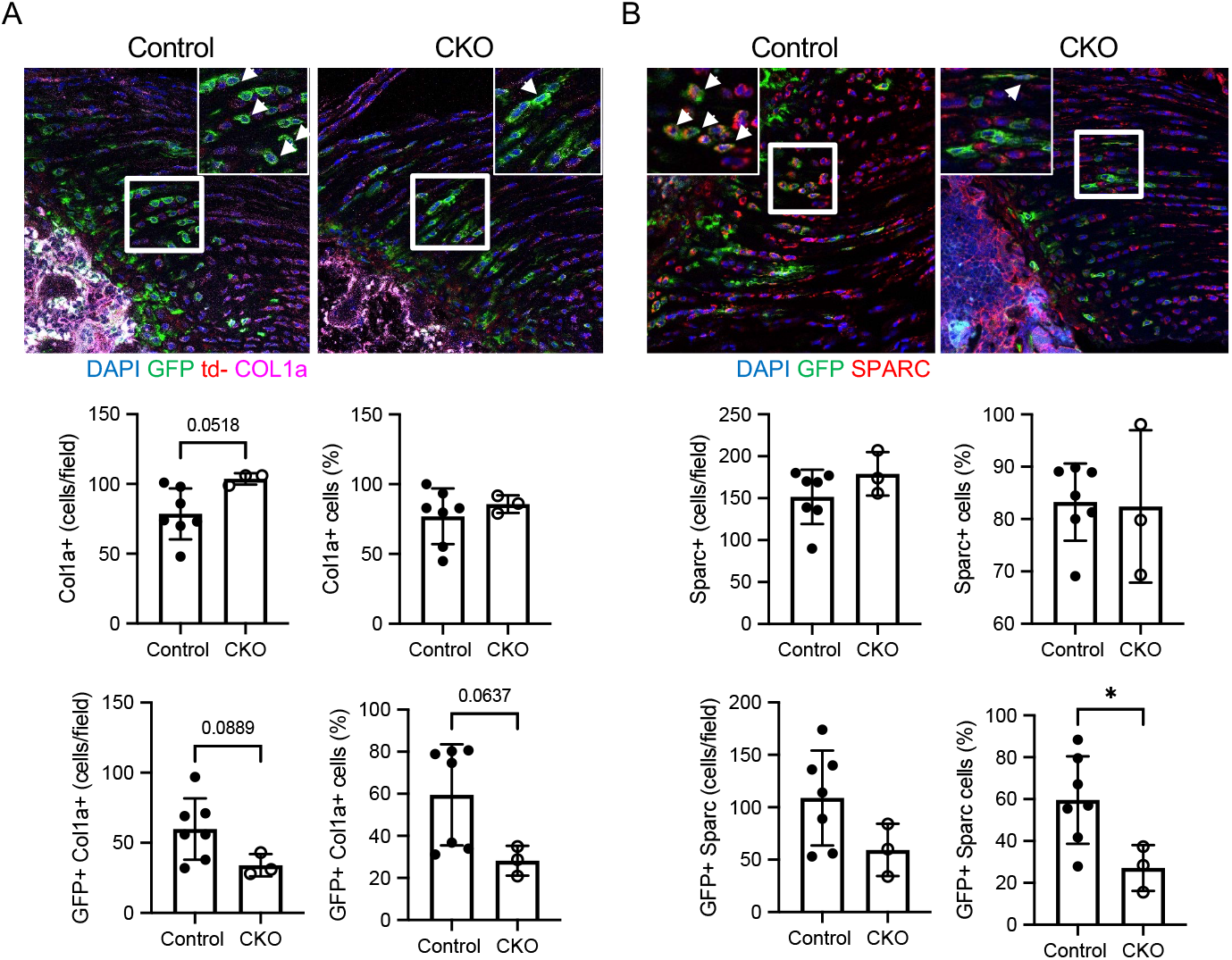
ECM changes in Gli1Cre^ERT2^;TbR2^fl/fl^ CKO mice were limited to Gli1-lineage cells. Immunohistochemistry for **(A)** COL1a (magenta) and **(B)** SPARC (red) in Control and CKO supraspinatus tendon entheses. Gli1-lineage cells are GFP+ (green) (bone is on the lower left and tendon is on the upper right of the sections). The insert is an enlarged view of the white boxed area of the enthesis (positive cells indicated by arrowheads). Histomophometric analyses demonstrated numbers and percentage of cells positive for (A) COL1a and (B) SPARC for the enthesis and for Gli1-lineage cells (i.e., colocalization of GFP with either COL1a or SPARC). * p<0.05.

### TbR2 deletion led to fewer Gli1-lineage cells and reduced proliferation

To see if TbR2 deletion in Gli1-lineage cells affected their proliferation, CTRL and CKO cells from tail tendon were cultured to passage 3 and analyzed using FACS. The number of GFP+ Gli1-lineage cells was significantly reduced in CKO mice compared to CTRL mice (Figure 4A). Similar results were seen for enthesis cells isolated from supraspinatus and infraspinatus tendon entheses (Figure S2). Consistent with these *in vitro* results, immunohistochemical analysis of mouse supraspinatus entheses for the proliferation marker Ki67 showed a trend of decreased Ki67+ cells in CKO entheses (Figure 4B, S1D). These data suggest that Gli1-lineage cell proliferation is in part dictated by TGFβ signaling.

**Figure 4.**
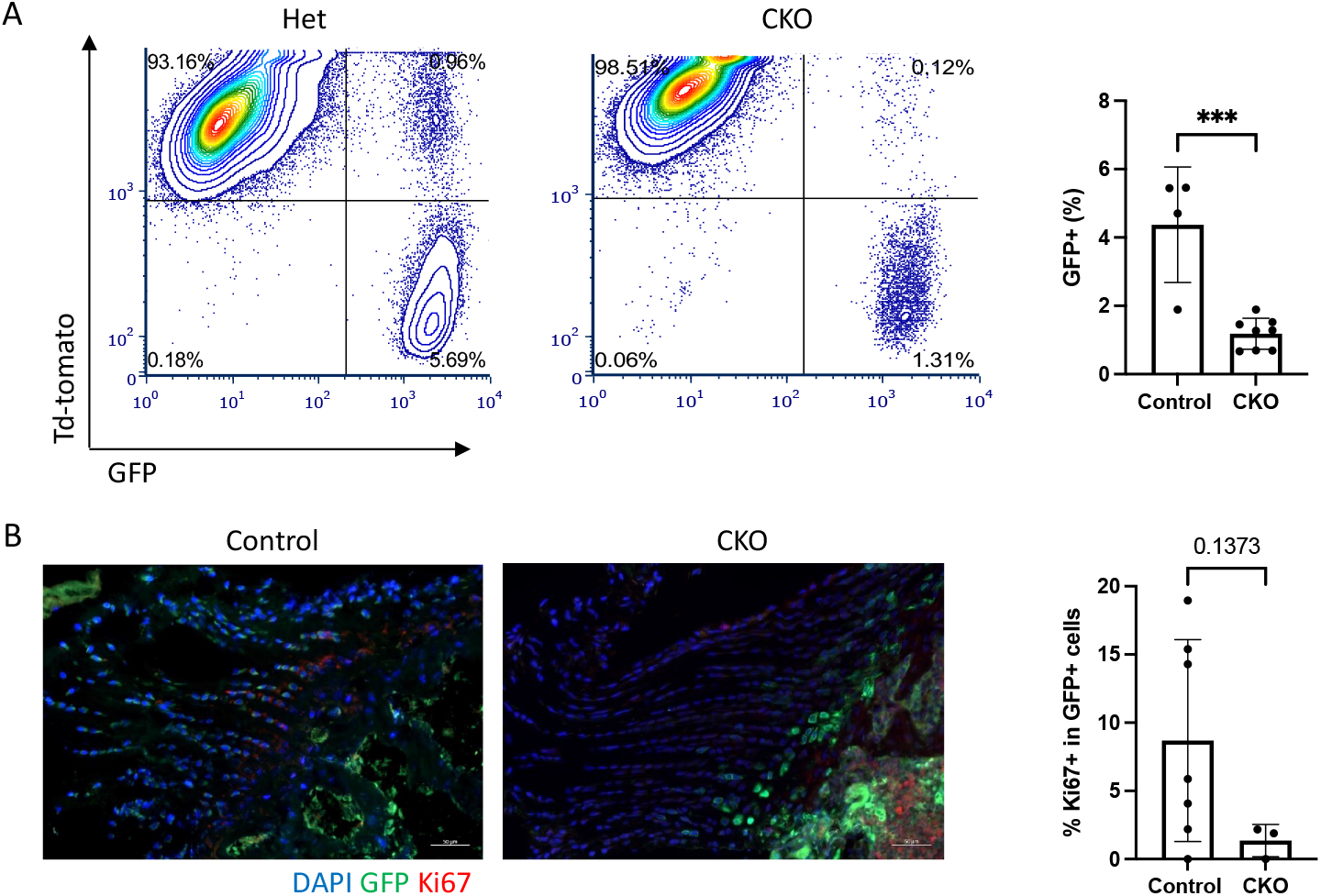
TbR2 deletion led to fewer Gli1-lineage cells and reduced proliferation. **(A)** FACS dot plots (left) and quantification (right) showed reduced numbers of Gli1-lineage cells in CKO mice. **(B)** Ki67 immunostaining showed decreased Ki67+ in Gli-lineage cells in CKO supraspinatus entheses (*** p<0.001; tendon is on the left and bone is on the right of the sections).

### Smad-dependent canonical and Smad-independent non-canonical pathways were affected in TbR2-deficient mice

FACS-sorted Gli1-lineage cells derived from tail tendon from Gli1CreERT2;TbR2^fl/wt^;mTmG control and Gli1CreERT2;TbR2 ^fl/fl^;mTmG CKO mouse TFs were stimulated with TGFβ 1 or hedgehog agonist (HhAg). TGFβ treatment increased Scx gene expression in a time-dependent manner in control cells, and this induction was significantly reduced in CKO cells (Figure 5). Similarly, HhAg induced Scx expression in a time dependent manner, but there was no difference between responses in the control and CKO cells (Figure 5). Sox9 was not significantly induced by either TGFβ or HhAg (Figure 5). As expected TGFβ induced expression of TGFβ 1 (Figure S3) and HhAg induced expression of Gli1 (Figure S4).

**Figure 5.**
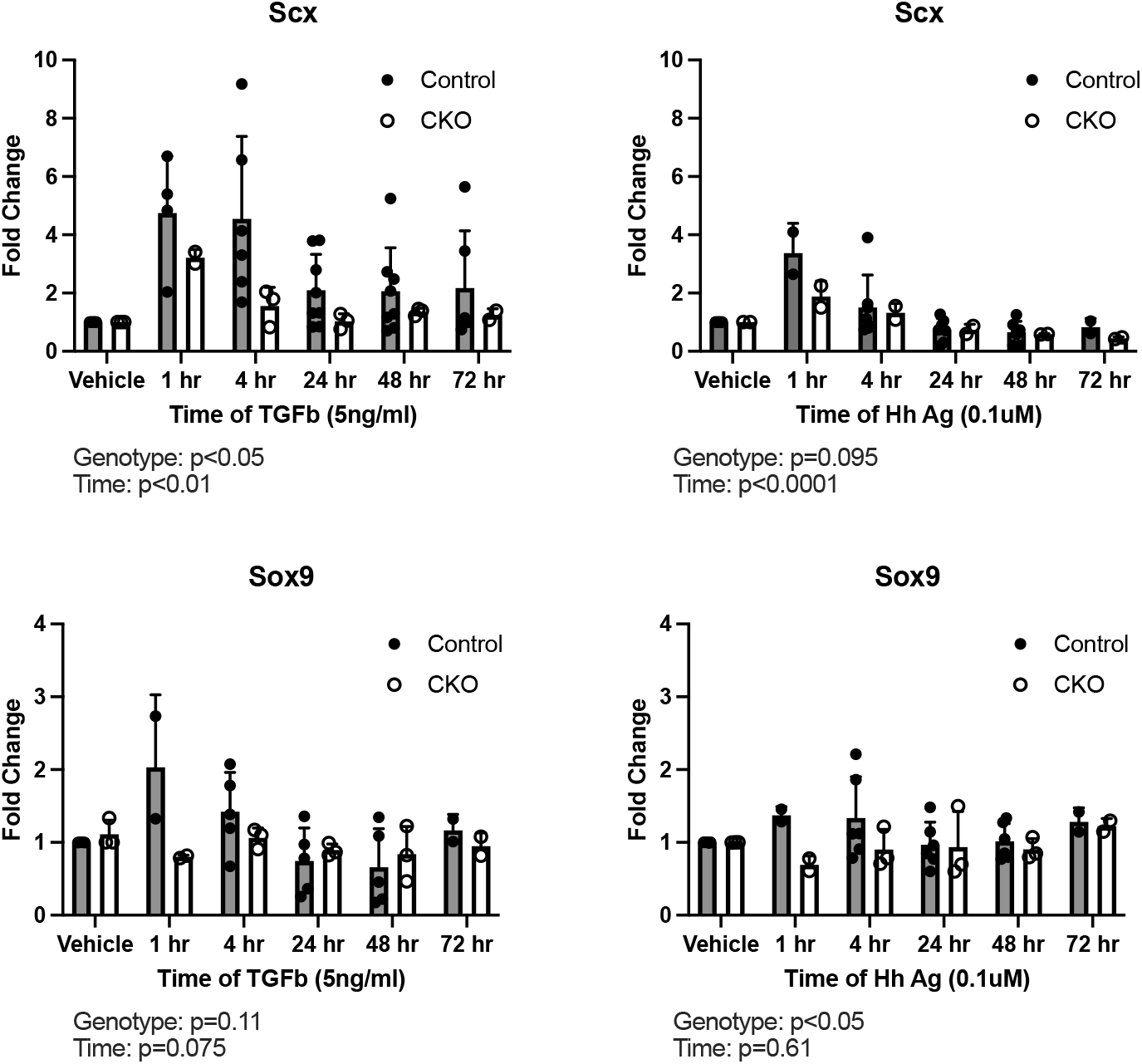
Responses of Gl1-lineage cells to TGFβ and HhAg *in vitro*. TbR2 deletion impaired TGFβ-induced Scx expression and HhAg-induced Sox9 expression in Gli1-lineage cultured cells. The effect of genotype and time was determined by two factor ANOVA.

To investigate whether Smad-dependent canonical or Smad-independent non-canonical pathways were affected by TbR2 deletion, Gli1-lineage cells were FACS sorted and stimulated with TGFβ. Whole cell lysates were probed for phosphorylated Smad2, the key downstream signaling molecule for the canonical TGFβ pathway, as well as phosphorylated cJun, JNK, p38, and Erk, involved in non-canonical pathways. As expected, Smad2 phosphorylation was significantly decreased in CKO cells (Figure 1C), suggesting that the canonical pathway was affected by TbR2 deletion. Both cJun and Erk were induced by TGFβ in a time dependent manner, but only cJun and JNK activation were TbR2-dependent, showing reduced phosphorylation in CKO cells (Figure 6). cJun activation was verified on tissue section using immunohistochemistry for the phosphorylated form of the cJun protein. The numbers and percentage of cJun+ cells were similar in CKO and Control entheses (Figure 6C). When considering only Gli1-lineage (i.e., GFP+) cells, there was a dramatic reduction in the percentage of cJun+ cells in CKO compared to control enthesis (Figure 6C). Notably, non-conical TGFβ signaling through MAPK/JNK/cJun pathways may controls cell proliferation [37-39], consistent with the result that TbR2 deletion led to fewer Gli1-lineage cells and reduced proliferation (Figure 4). Akt activation was not induced by TGFβ (data not shown). These results suggest that both Smad2 canonical and cJun non-canonical pathways were dependent on TbR2 in Gli1-lineage cells.

**Figure 6.**
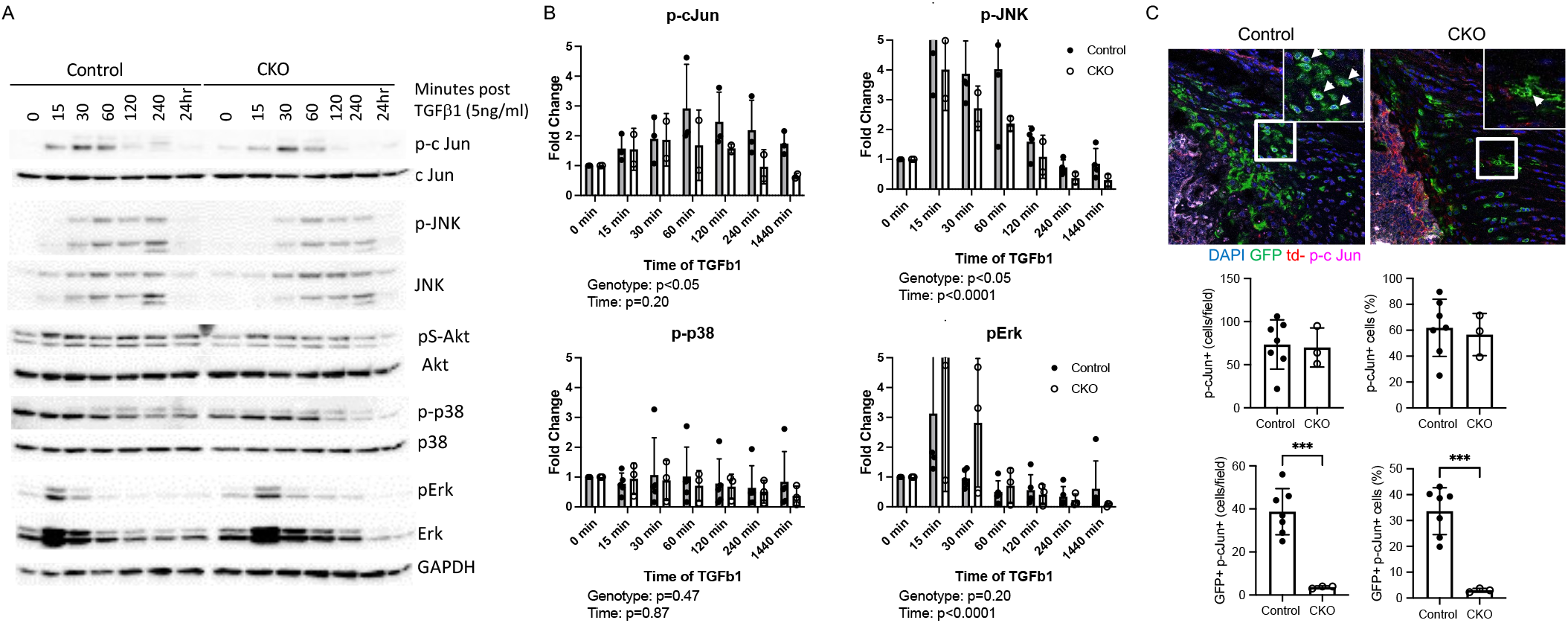
TGFβ induced non-canonical pathways in Gli1-lineage cells. **(A)** Representative western blots for control and CKO Gli1-lineage cells. **(B)** Quantification of phosphorylated cJun, JNK, p38, and Erk in control and CKO Gli1-lineage cells. The effect of genotype and time was determined by two factor ANOVA. **(C)** Immunofluorescent staining of p-cJun (magenta) in Control and CKO supraspinatus tendon entheses. Gli1-lineage cells are GFP+ (green) (bone is on the lower left and tendon is on the upper right of the sections; positive cells indicated by arrowheads).

### TGFβ signaling drove Scx expression through its distant enhancer

Previous studies have shown that TGFβ drives Scx gene expression in tendon cells [20]. However, this has not been explored in Gli1-lineage cells and the molecular mechanism by which TGFβ regulates Scx remains elusive. PROMO software was used to search 10 kilobases (kb) up- and down-stream of the mouse Scx gene transcription start site (TSS), revealing three potential Smad4 binding sites when the dissimilarity parameter was set at 4.0 (Figure 7A). Using ChIP with an anti-Smad4 antibody, the strongest binding site was located ∼4.9 kb downstream of the Scx TSS (Figure 8B). Binding of Smad4 to this site depended on TbR2, as TGFβ -induced binding was diminished in TbR2 deficient cells (Figure 7C). These data provide direct evidence that TGFβ controls Scx through Smad4 binding to the Scx enhancer 4.9 kb downstream from its transcription starting site in those cells.

**Figure 7.**
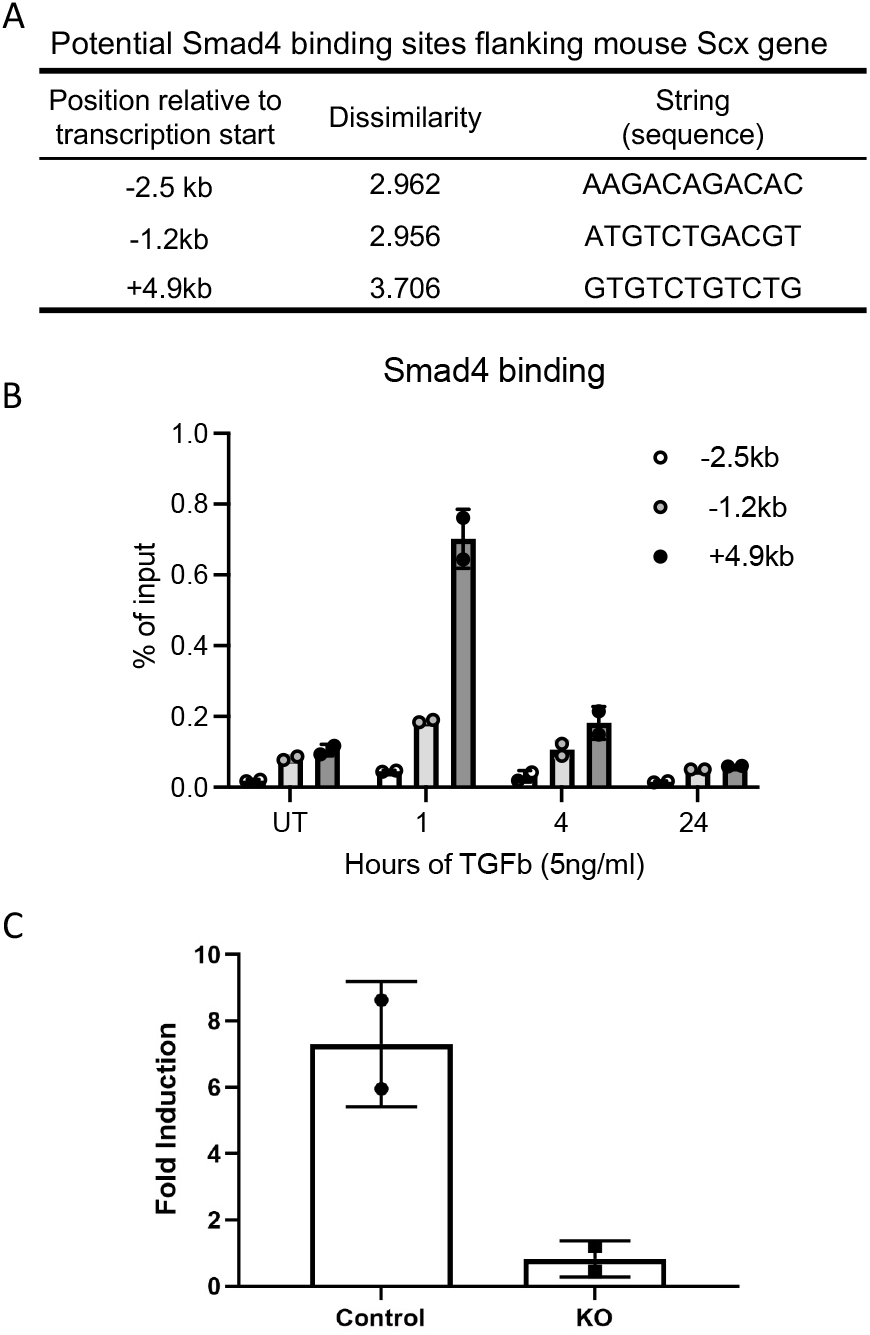
TGFβ controls Scx expression through Smad4 binding to a distant Scx enhancer. **(A)** Potential Smad4 binding sites in the mouse Scx locus. **(B)** Time-dependent Smad4 binding to 3 potential sites. **(C)** Smad4 binding to 4.9 kb downstream of the Scx enhancer was TbR2-dependent.

## DISCUSSION

The enthesis is a unique mineralized fibrocartilage tissue that bridges tendon and bone [40, 41]. This functionally graded transitional tissue is formed by a pool of Gli1+ (i.e., hedgehog responsive) cells that eventually populate the entire enthesis [12, 23]. The TGFβ family of growth factors have been shown to play important roles in the development, maintenance, and healing of musculoskeletal tissues, including bone, cartilage, and muscle [42, 43]. TGFβ promotes bone growth by enhancing hypertrophic chondrocyte differentiation and calcification of the bone collar through Runx2 and Wnt pathways, as well as cilia [18, 44-48]. TGFβ signaling has also been shown to drive the formation of tendon and the enthesis; loss of TGFβ signaling results in defective tendon and ligament formation [20, 21] and abnormal enthesis mineralization [49]. Furthermore, recent reports have suggested that TGFβ controls fibrocartilage and ligament cell proliferation [22, 50]. However, the molecular mechanisms by which TGFβ controls tendon and enthesis formation remain unclear. Therefore, the current study used an inducible conditional knockout model to delete TbR2 in Gli1-lineage cells postnatally. We hypothesized that deletion of TbR2 in Gli1-lineage enthesis cells would negatively affect development and mineralization of the enthesis. Since enthesis mineralization occurs at approximately P14 [51, 52], we induced TbR2 deletion at P5-P7 [23]. Loss of TGFβ signaling at the postnatal enthesis led to defects in humeral head bone morphology and enthesis mechanical properties by P56 and reduction of ECM proteins COL1a and SPARC in Gli1-lineage cells. This is consistent with prior studies showing the importance of the TGFβ superfamily for tendon and enthesis development [20, 53].

To explore mechanisms by which TGFβ regulates tendon enthesis cell differentiation and proliferation, a mouse reporter model was developed. The model was based on the important role that Gli1-lineage cells play in the development of a variety of tissues, including the enthesis [23]. Isolating these cells is a crucial step to understand their function and molecular mechanisms. Using Gli1Cre^ERT2^;mTmG reporter mice combined with flow cytometry, we successfully isolated these cells to a high level of purity from mouse rotator cuff tendon entheses and from tail tendons. Specifically, Gli1Cre^ERT2^;TbR2^fl/fl^;mTmG mice allowed for the isolation of GFP-expressing Gli1-lineage cells that had TbR2 deleted and td-Tomato expressing cells that did not express Gli1 and had TbR2 intact. This tool allowed us to explore TGFβ regulation of Gli1+ cells both *in vitro* and *in vivo*.

At the cellular level, TGFβ signaling is initiated by binding of ligands such as TGFβ 1, β 2, and β 3, leading to dimerization of TbR2 with TbR1 and downstream phosphorylation of Smad2 and 3. Phosphorylated Smad2/3 then further phosphorylates and combines with Smad4. Phosphorylated Smad4 brings other Smad proteins to the nucleus and binds to DNA to promote gene transcription leading to cell proliferation, differentiation, and/or ECM deposition [54, 55]. Other than TGFβ /Smad canonical signaling pathway, TGFβ also activates non-canonical pathways through TAK/MAPK and mTOR, which controls cell proliferation and differentiation [42, 56, 57]. Both *in vitro* and *in vivo*, the JNK/cJun non-canonical pathway showed TbR2 dependency in Gli1-lineage cells, with marked reduction in cJun activation in CKO enthesis Gli1-lineage cells. These results coincided well a reduction in total and proliferating Gli1-lineage cells in CKO mice and imply that JNK/cJun-mediated non-conical TGFβ signaling is necessary for proliferation of Gli1-lineage cells.

At the molecular level, TGFβ has been shown to upregulate key transcription factors for tendon and enthesis development (e.g., Scx, Mohawk) [20, 58, 59]. Furthermore, it was reported that Smad3 could physically associate with Scx and Mohawk and regulate their expression [60]. Using Gli1-expressing cells isolated from our reporter mouse model, with or without TbR2, we found that the expression of Scx was increased by TGFβ in control cells, whereas this induction was blocked in TbR2-deleted Gli1+ cells. It has been shown that Smad proteins directly associate with Scx in a cell line [60]. However, the effect of this interaction is not clear. To explore how TGFβ controls Scx expression at the molecular level, we carried out a ChIP assay using an anti-Smad4 antibody, as Smad4 mediates all of the Smad protein DNA binding activities [61, 62]. ChIP results demonstrated that Smad4 binds to the enhancer of Scx at the +4.9kb position, and this binding was significantly diminished in TbR2-deleted cells. A recent study reported a similar binding motif for SMAD4 in CD8+ T cells [63]. Notably, this distal enhancer is conserved among mammals, and is thus highly likely to function similarly in other animals, including human [30]. Further mutagenesis studies are needed to prove this is the bona fide Smad4 binding site regulating Scx expression.

Using tools ranging from knockout animal to cellular and molecular biology, the results of the current study add to the growing body of literature defining the regulation of the Gli1-lineage cells that build and mineralize the enthesis. TGFβ signaling is necessary for the formation of a functional enthesis, affecting both differentiation and proliferation of Gli1-lineage enthesis cells, likely via canonical and non-canonical pathways, respectively. This work can be applied in future translational studies to drive the regeneration of the enthesis during tendon-to-bone healing via growth factor and/or stem cell therapies.

## ACKNOWLEDGEMENTS

The study was supported by the NIH/NIAMS through R01 AR057836. Flow cytometry analysis was supported by NIH grant S10 BR027050 and cell sorting was supported by NIH grant S10 OD020056. Microscopic core was supported by NIH grant P30 CA013696.

## Supplemental Document

**Figure S1.**
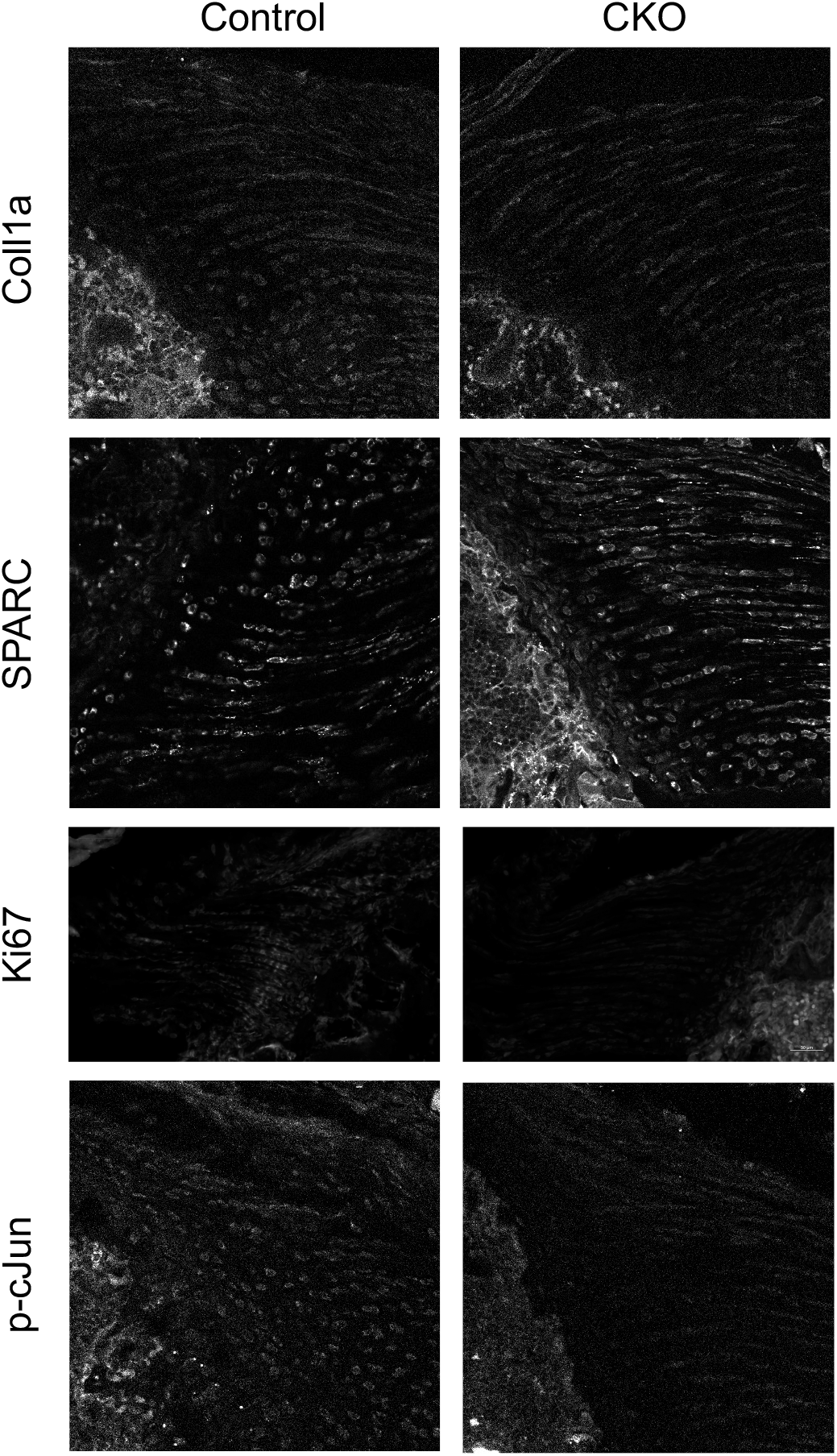
Immunohistochemistry for COL1a, SPARC, Ki67, and p-cJun in mouse supraspinatus tendon entheses. Single channel images demonstrating representative stainings for COL1a, SPARC, Ki67, and p-cJun (bone is on the lower left and tendon is on the upper right of the sections for COL1a, SPARC, and p-cJun; bone is on the right and tendon is on the left of the sections for Ki67).

**Figure S2.**
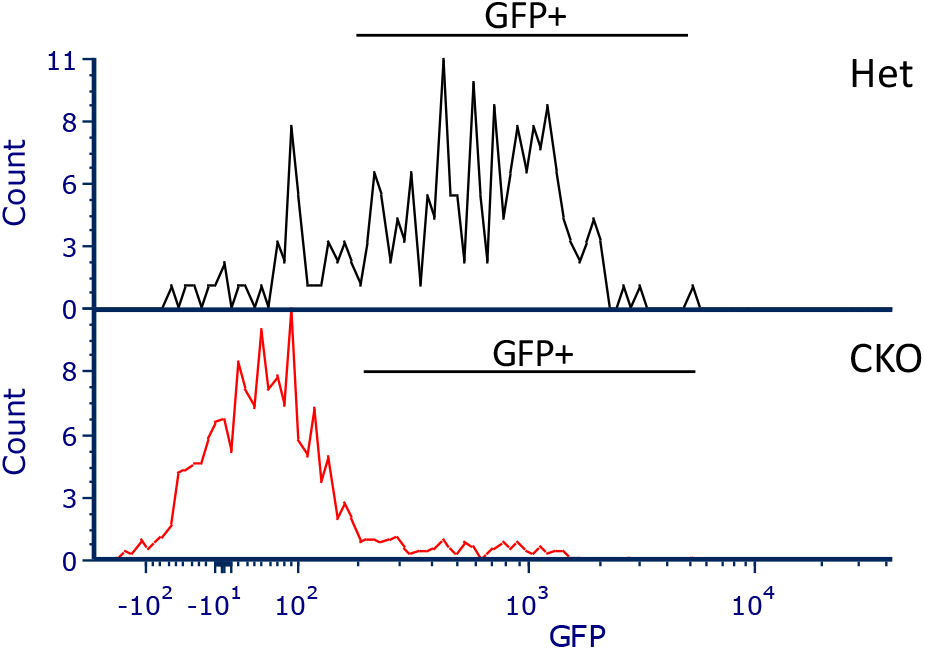
Histogram showing there were fewer GFP+ lineage cells in TbR2 CKO entheses compared to control entheses. The data were representative of two independent experiments.

**Figure S3.**
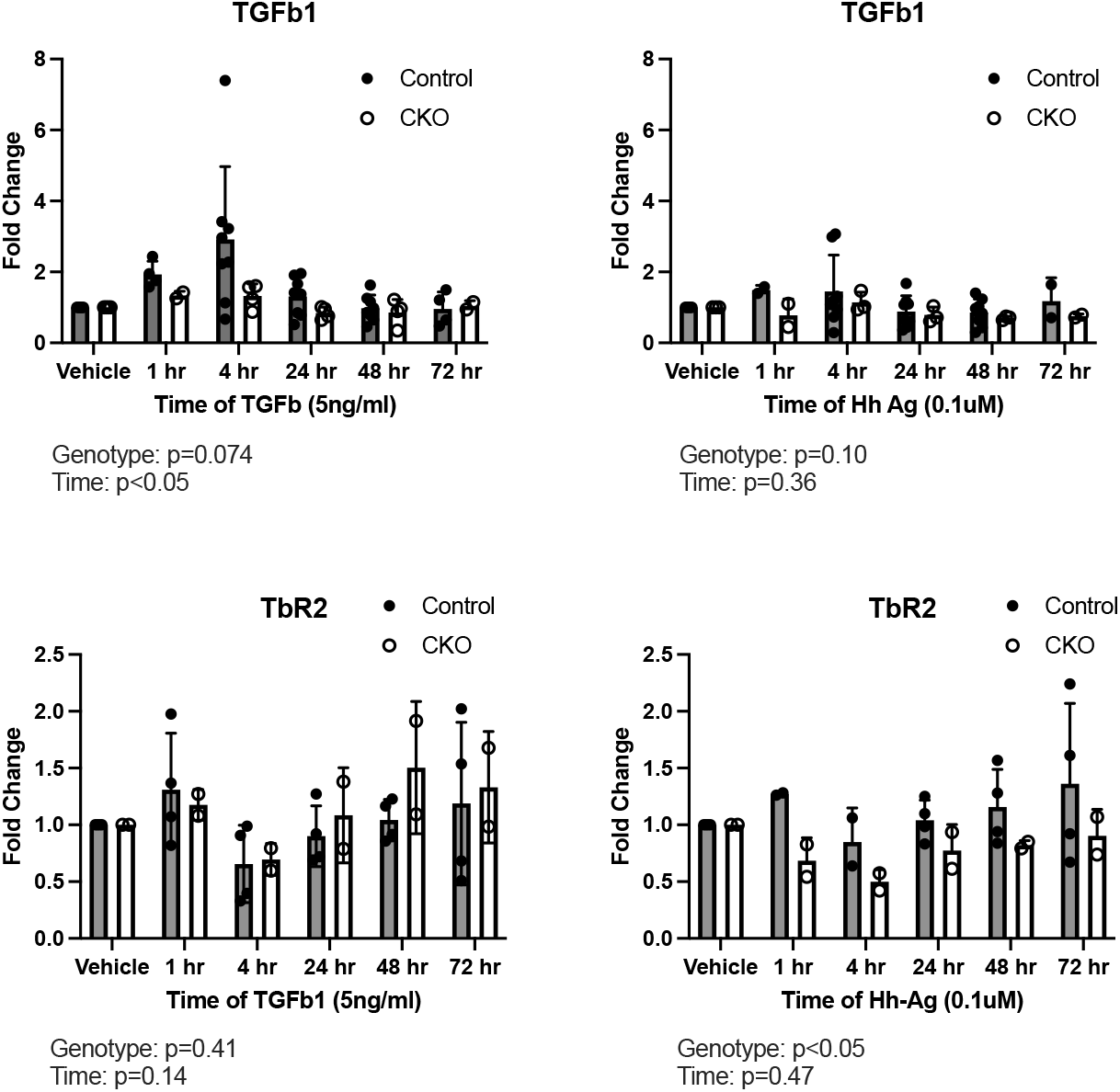
TGFβ -related responses of Gl1-lineage cells to TGFβ and HhAg *in vitro*.

**Figure S4.**
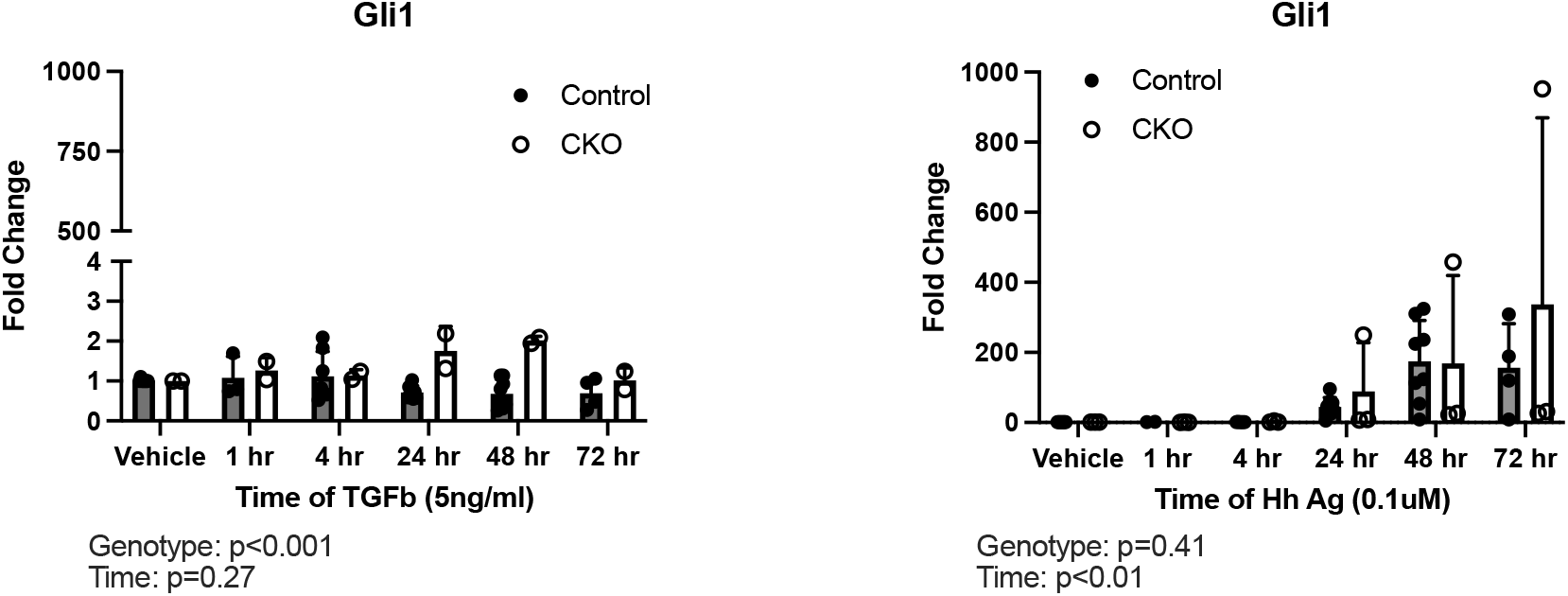
Hedgehog-related responses of Gl1-lineage cells to TGFβ and HhAg *in vitro*.

